# TNFR2 is Expressed by a Discrete Subset of Epidermal γδ T Cells with an IL-17 Gene Signature during Psoriasis

**DOI:** 10.1101/2025.09.05.674535

**Authors:** Jasmine Ahmed, Carolyn Matthews, Patricia Tulloch, William Lawler, Albert Mendoza, Mikaela Rhoiney, Julie M. Jameson

**Author notes:** Corresponding Author: Dr. Julie Jameson, Department of Biological Sciences, California State University San Marcos, 333 S. Twin Oaks Valley Rd, San Marcos, CA 92096. Authors contributed equally to this work (Jasmine Ahmed, Carolyn Matthews, Patricia Tulloch).

## Abstract

Psoriasis is a chronic skin disease that results in scaly patches and affects 2-3% of people worldwide. Therapeutic treatment targets the TNF-α/IL-17 axis to disrupt keratinocyte hyperproliferation and inflammation. While more is known about the role of dermal αß and γδ T cells in IL-17 production, less is understood about the role of resident epidermal T cells. Here, we examine how TNF-α modulates epidermal γδ T cell activation and function. We show that a subset of activated epidermal γδ T cells expresses TNFR2 with or without TNFR1. Stimulation with TNF-α induces epidermal γδ T cells to produce IL-17 family cytokines and chemokines. Epidermal γδ T cells do not require TNFR1 or 2 for development or homing to the skin. Instead, TNFR2 plays roles in epidermal γδ T cell function skewing them toward a Tγδ17 phenotype during psoriasis. Investigation of the mechanisms by which TNF-α associated inflammation impacts epidermal γδ T cell function may help identify new cellular targets for immunotherapy or mark them as early regulators of skin inflammation.

## Introduction

Psoriasis is a chronic inflammatory skin disease that affects approximately 7.5 million adults in the United States (1,2). Genetics, the environment, and the immune system are major contributors to psoriasis disease pathogenesis (3–5). Patients with the most common form of psoriasis, psoriasis vulgaris, exhibit scaly patches on their skin characterized by epidermal hyperplasia, leukocyte infiltration, and dysregulated cytokine production. The TNF-α/IL-23/IL-17 cytokine axis is associated with psoriasis and a target for therapeutic treatment (6). The cellular mechanism involves complex crosstalk between a variety of cells including keratinocytes and dendritic cells which respond to stress, damage, or infection in the skin through the production of Type I IFNs, IL-23 and TNF-α (7,8). These cytokines either directly or indirectly induce IL-17 production by pathogenic αβ and γδ T cells that migrate to the epidermis (9,10). IL-17 induces keratinocyte proliferation and can synergize with TNF-α to further increase psoriatic gene expression (11–14). TNF-α regulates T cell proliferation in murine models of psoriasis and inhibition of TNF-α in patients improves clinical outcomes (15,16). While the focus has been on infiltrating αβ and γδ T cells, recent findings have identified a resident IL-17-producing epidermal γδ T cell population in psoriasis (17).

Skin γδ T cells play key roles in wound healing and skin homeostasis in mice and humans (18– 20). Epidermal T cells in mice express a canonical Vγ5Vδ1 TCR (Heilig and Tonegawa nomenclature (21)), which recognizes damaged, stressed, or transformed keratinocytes (22,23). Activated epidermal γδ T cells produce cytokines to modulate inflammation, release growth factors to promote wound reepithelialization, and perform cytolysis to eliminate tumors (18,24,25). These diverse functions suggest that either epidermal γδ T cells exhibit plasticity, or there are distinct functional subsets. Epidermal γδ T cell function is regulated through costimulation, but less is known about how cytokine reception may regulate functional plasticity or subset differentiation and activation (26,27).

Transmembrane and secreted TNF-α acts through two receptors, TNFR1 and TNFR2 to promote pleiotropic effects from proliferation to cell death. TNFR1 is widely expressed and TNF-α binding initiates NF-kB and MAP kinase signals along with caspases (28). TNFR2 expression is restricted to immune cells, neural cells, and endothelial cells where it is involved in both cellular proliferation, apoptosis, and cytokine production. In murine models of psoriasis induced by imiquimod (IMQ), the role of TNFR1 and 2 are complex with some studies showing that mice deficient in TNFR1 exhibit reduced inflammation and mice deficient in TNFR2 show no effect (29). While other studies show that TNFR2 deficient mice exhibit reduced inflammation in the IMQ psoriasis model (30). Still others show TNFR2 knockout mice exhibit more severe disease (29). These conflicting data attest to the complexity in cellular TNFR expression, resulting function, and alterations in chronic inflammation. Thus, in psoriasis, it is important to elucidate how resident epidermal T cells are modulated by the TNF-α/IL-23/IL-17 cytokine axis.

Here, we examine how TNF-α impacts epidermal γδ T cell activation and function to determine how TNFR1 and/or TNFR2 modulate functional outcome. We show that a subset of epidermal γδ T cells express TNFR2 and that TNF-α induces the production of a specific set of cytokines. While epidermal γδ T cells do not require TNFR1 or 2 for development or homing to the skin, TNFR2^+^ epidermal γδ T cells express Tγδ17 signature genes during psoriasis. Investigation of the mechanisms by which TNF-α associated inflammation impacts epidermal γδ T cell function may help identify new cellular targets for immunotherapy or mark them as early regulators of skin inflammation.

## Materials and Methods

### Mice

Male C57BL/6J (B6) +/+ and B6.129S-Tnfrsf1a ^tm1/mx^ (TNFR1-/-) and Tnfrsf1b ^tm1/mx^ (TNFR2-/-) mice were purchased from the Jackson Laboratory (Bar Harbor, ME). B6.129S-Tnfrsf1a ^tm1/mx^ and Tnfrsf1b ^tm1/mx^ were backcrossed six generations and crossbred to generate double knockout (DKO) B6 TNFR1-/-TNFR2-/-mice. Male C57BL/6NTac were purchased from Taconic Biosciences, Inc. (Hudson, NY, USA). Mice were used between the ages of 8 – 16 weeks for cell line generation. For diet-induced obesity, 6-week-old mice were fed a high fat diet consisting of 60 kcal% from fat (HFD) (D12492; Research Diets, Brunswick, NJ) or a normal chow diet consisting of 17 kcal% from fat (NCD) (7012; Harlan Envigo) for 27-33 weeks. Mice were housed in specific pathogen-free conditions at California State University San Marcos according to California State University San Marcos IACUC guidelines (nos. 18-007, 21-003).

### Epidermal T Cell Isolation and Culture

Epidermal cells were isolated from wildtype B6 mice as previously described (18,31). Briefly, skin from the back was excised and cut into 1-cm^2^ squares. Skin was incubated on 0.3% trypsin-GNK (.09% glucose, .84% sodium chloride, and .04% potassium chloride) at 37°C with 5% CO_2_ for 3 hours. Epidermal sheets were peeled from the dermis and shaken in 0.3% trypsin-GNK with .1% DNase at 37°C for 10 minutes. The cells were filtered through sera separa filter columns (Evergreen Scientific) and isolated with a Lympholyte M gradient (Cedarlane Labs). Cells were cultured in 10% RPMI with 2.0 ug/mL Con A, 1.0 ug/mL indomethacin, and 20 U/ml IL-2 (10% fetal bovine serum, 2.5% HEPES buffer, 1% nonessential amino acids, 1% sodium pyruvate, 1% penicillin-streptomycin-glutamine, .1% 2-mercaptoethanol, and 20 U/ml IL-2). Media was replaced every 3-4 days with 10% RPMI with 20 U/mL IL-2. Cells were re-stimulated with 1.0 μg/mL of Con A every 3 weeks.

### Epidermal T cell Activation and Flow Cytometry

After 5-12 weeks of culture, cells were harvested and activated with 1μg/ml anti-CD3 (Biolegend) with or without low (10 ng/mL) or high (100 ng/mL) recombinant mouse TNF-α (Biolegend) for 24 hours. Cells were stained with antibodies specific for CD3e (clone 145-2C11), TNFR2 (CD120b), CD25 (PC61), TNFR1 (CD120a) (Biolegend), TCR Vγ5 (clone 536) (BD Biosciences). Flow Cytometry was performed on a BD Accuri™ C6 Flow Cytometer (BD Biosciences) and analyzed with FlowJo v10 software (BD Biosciences). For flow cytometry plots, gating was determined for each individual experiment using negative or isotype controls.

### Epidermal Sheet Preparation and Immunofluorescence Microscopy

Epidermal sheets were isolated and stained as previously described (18,31). Briefly, ears were split in half and placed on 3.6% ammonium thiocyanate in PBS for 10-15 minutes. Epidermal sheets were peeled from the dermis and stained with antibodies specific for TNFR2 and TCR Vγ5 for 1 hour at 37°C with 5% CO2. Epidermal sheets were mounted using Slowfade Gold Antifade Reagent with DAPI (Invitrogen). Digital images were acquired (Nikon Microphot-FXL) at original magnification x 200 and analyzed using Adobe PhotoShop CS5 software (Adobe Systems Incorporated, San Jose, CA, USA). Cells were quantified using grids and calculated as cells/mm^2^. At least three separate experiments were performed and a minimum of 500 cells were quantified per experiment.

### Luminex Multiplex Assay

Supernatants harvested from activated epidermal γδ T cells were analyzed using the Cytokine/Chemokine 26-Plex Mouse ProcartaPlex (EPXR260-26088-901, Thermo Fisher) on the Luminex 200 instrument (Thermo Fisher). Briefly, a series of standards were run in duplicate and cell culture supernatants were run in triplicate to detect cytokines and chemokines. Data was analyzed using xPONENT 3.0 software (Diasorin).

### Single cell RNA Sequencing (scRNAseq) Data Analysis

scRNAseq data from Liu et al. was downloaded as fastq files from the NIH Gene Expression Omnibus (GEO) database (GSE149121) (32). Files were downloaded to the Linux terminal and transferred to the Cellranger Analysis Pipeline terminal. All fastq files were individually run through Cellranger Analysis Pipelines v6.1 (10x Genomics) to count the cells. After running the cell count, the fastq files were run through Cellranger again to aggregate and structure the files to compare IMQ-treated murine cell sets against the untreated control groups. All counted cells were aggregated alongside the mm10 murine genome for reference and consistency. Once aggregation and structuring of the data was completed, Cellranger generated a cloupe file that was uploaded to the visualization software Loupe Browser v6.0 (10x Genomics) to be used for downstream analysis of scRNA seq data. Cells positive for expression of *CD3, Tcrg-v5*, and *Fcer1g* were identified as epidermal γδ T cells. Any cells positive for *Tcrg-v4, Tcrg-v6, Cd4*, or *Cd8* were sorted out as non-target cells. Once the epidermal γδ T cell cluster was identified, further analysis was performed by clustering the epidermal γδ T cells into subsets based on their expression of Tnfrsf1a (TNFR1) and Tnfrsf1b (TNFR2). Elucidated subsets were compared between IMQ-treated and untreated controls to establish differential gene expression between epidermal γδ T cell subsets with and without psoriasis-like disease. The differentially expressed genes between these subsets were exported from Loupe Browser and submitted to Ingenuity Pathway Analysis (Qiagen) for core analysis.

### Statistical Analysis

Data was analyzed using GraphPad Prism software (La Jolla, CA, USA) and represented as mean ± standard error of the mean (SEM). Two-way repeated measures analysis of variance (RM-ANOVA) with Bonferroni multiple comparisons and Student’s t-test were utilized to compare multiple groups with two factors or two groups, respectively. All findings are considered significant at P < 0.05. Log2-fold change in Loupe Browser was calculated by using the localized ratio of normalized mean gene unique molecular identifier counts in each cluster relative to all other clusters.

## Results

### A subset of epidermal γδ T cells upregulate TNFR2 upon activation

To examine TNFR1 and TNFR2 expression, epidermal γδ T cells were stimulated with and without anti-CD3 in the presence or absence of TNF-α at low (10ng/ml) and high (100ng/ml) concentrations and analyzed by flow cytometry. Live, γδ TCR high cells were gated as these represent epidermal γδ T cells (Fig. 1A, 1D). Gated γδ T cells do not express TNFR1 or 2 prior to stimulation (Fig. 1A-C). TNF-α stimulation alone does not increase TNFR1 expression and only impacts TNFR2 expression at high concentrations (.97% to 16.8%) (Fig. 1B), but this increase does not reach statistical significance (Fig. 1C). Activation of epidermal γδ T cells with anti-CD3 increases TNFR2 expression by a population of epidermal γδ T cells (Fig. 1B, C). In contrast, there is not a significant increase in TNFR1 expression upon activation or TNF-α treatment (Fig. 1B).

**Figure 1.**
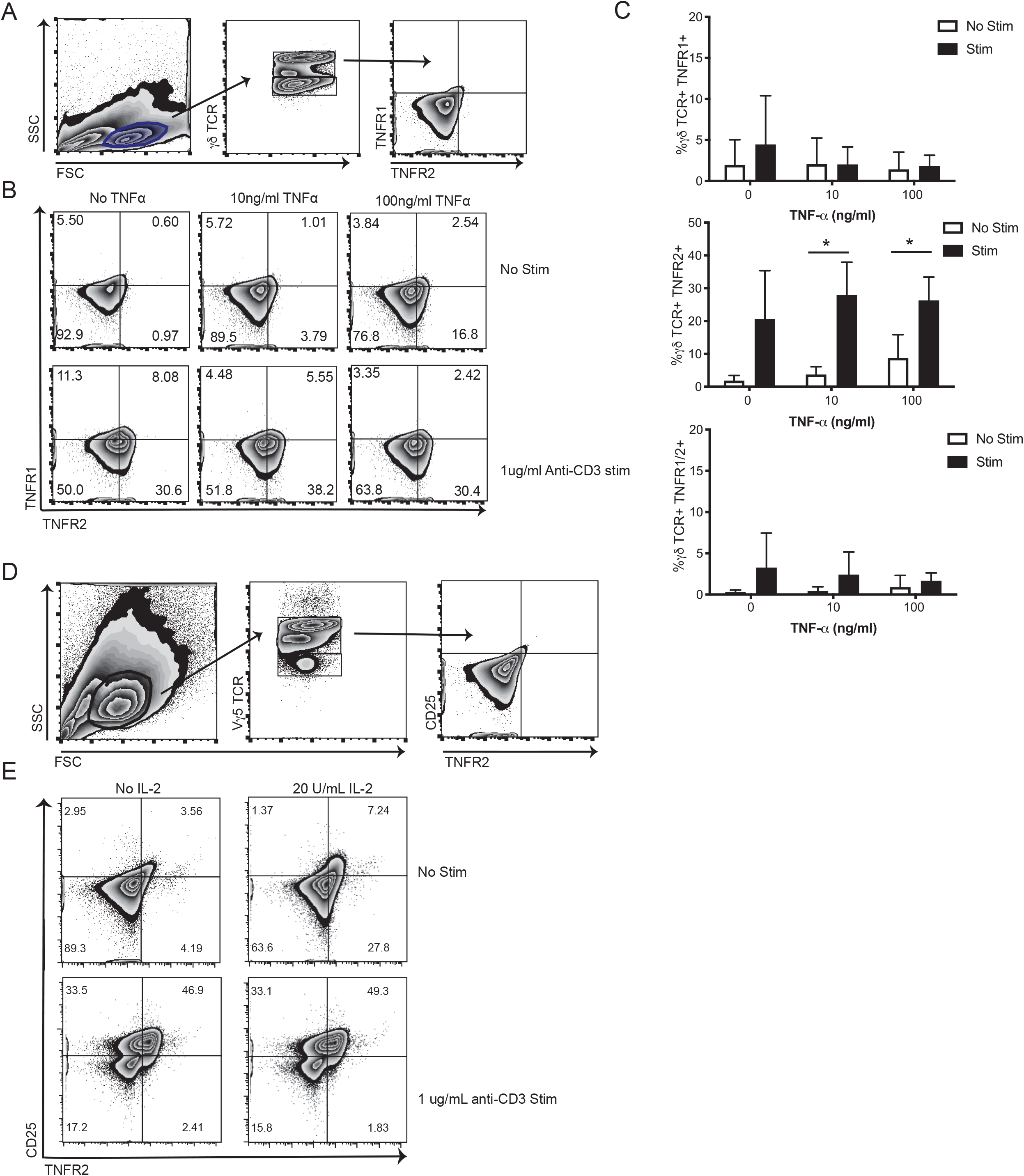
A subset of activated epidermal γδ T cells upregulates TNFR2 expression. (A) Flow cytometric analysis of epidermal γδ T cells. (B, C) Flow cytometry plots depicting live, γδ TCR^+^ cells expressing TNFR1 and TNFR2 after anti-CD3 stimulation with or without TNF-α treatment for 24 hours. n=3 (D, E) Flow cytometric analysis of epidermal VΨ5^+^ T cells after anti-CD3 stimulation with or without IL-2 treatment. n=2 *p < 0.05

We stained epidermal T cells with a Vγ5-specific antibody to further verify that the gated cells upregulating TNFR2 are epidermal γδ T cells. A population of live, epidermal Vγ5 T cells upregulate TNFR2 in the presence of anti-CD3 (Fig. 1D, E). In addition, almost all TNFR2-expressing Vγ5 T cells also express CD25 suggesting a role for IL-2 reception for this subset (Fig. 1E). Further, IL-2 alone induces some TNFR2 expression by Vγ5 T cells. This data identifies three main subsets of activated epidermal γδ T cells (1) TNFR2+CD25+ (2) TNFR2-CD25+ (3) TNFR2-CD25-.

### Epidermal γδ T cells do not require TNFR1 or 2 for seeding in the skin or maintaining homeostatic numbers

To determine the impact of TNFR1 and 2 on epidermal γδ T cell development, seeding and homeostasis, we examined epidermal γδ T cell numbers in mice lacking both TNFR1 and TNFR2. γδ T cells develop and seed the epidermis in TNFR1-/-TNFR2-/-(Fig. 2A). The number of epidermal γδ T cells in TNFR1-/-TNFR2-/- is similar to wild type mice (Fig. 2B). Thus, epidermal γδ T cells do not require TNFR1/2 for development and seeding in the skin. Further, the data also shows that TNFRs are not required by epidermal γδ T cells for homeostatic maintenance. To understand if epidermal γδ T cells express TNFR2 *in vivo*, epidermal sheets were stained with antibodies specific for Vγ5 and TNFR2 (Fig. 2C). TNFR2 expression is found on a small number of Vγ5^+^ T cells in the epidermis of wild type mice (Fig. 2C, D). We further examined if similar numbers of TNFR2-expressing Vγ5^+^ cells would be found in obese mice which are known to have chronically elevated TNF-α levels. Similar numbers of TNFR2^+^ Vγ5 T cells were observed in lean and obese mice suggesting that chronic TNF-α does not impact numbers of TNFR2-expressing T cells *in vivo*.

**Figure 2.**
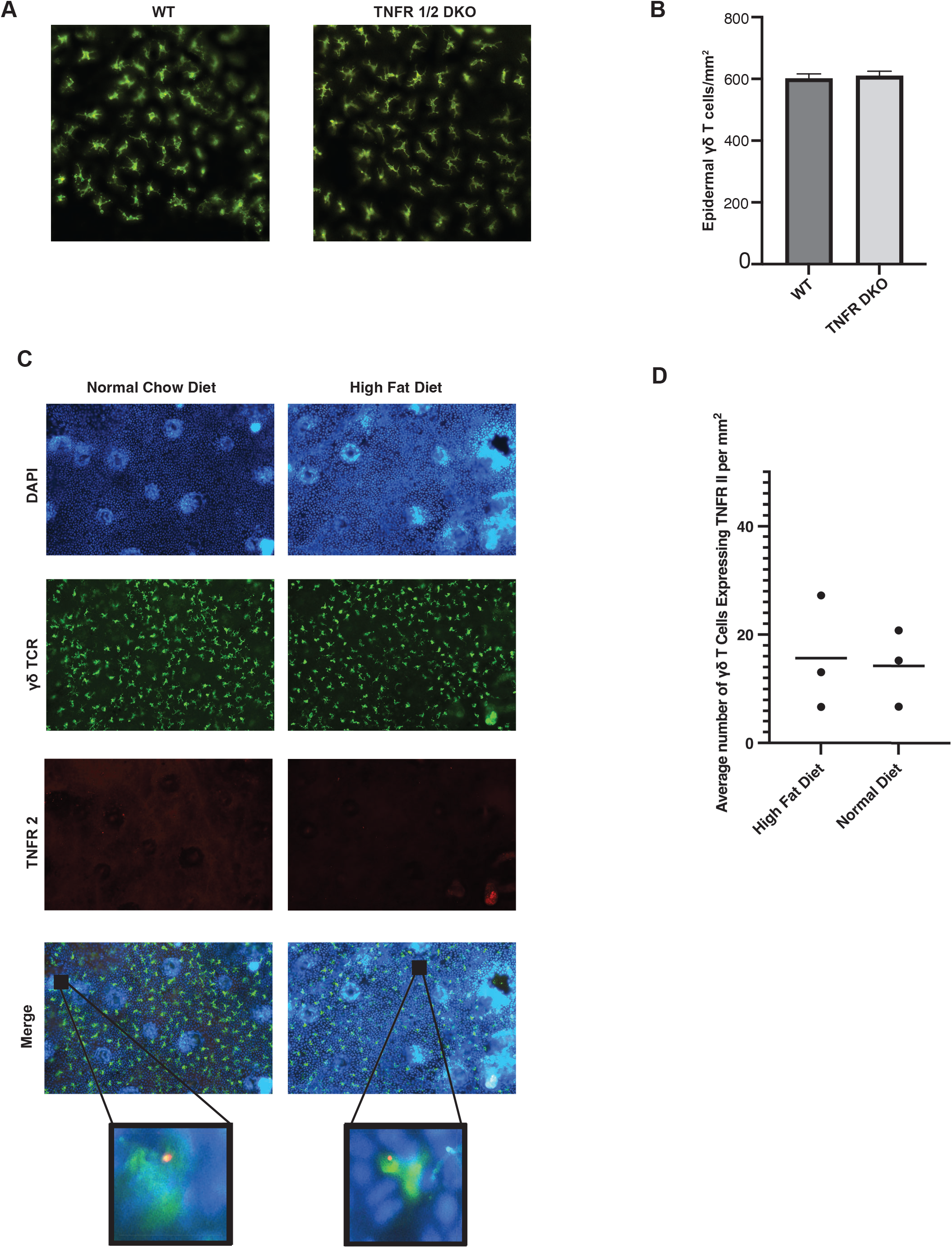
γδ T cells do not require TNFR1/2 for seeding and maintenance in the epidermis. (A, B) Representative images and quantification of epidermal sheets showing γδ T cells in WT and TNFR1/2 DKO mice (γδ TCR, green). n = 3 (C, D) Representative image and quantification of epidermal sheets from WT mice fed normal chow or HFD (γδ TCR, green, TNFR2, red). n=3 *p < 0.05

### While epidermal γδ T cells produce Th1, Th2, Th17 cytokines and chemokines upon anti-CD3 stimulation, TNF-a stimulation alone induces a Th17 focused response

To determine which cytokines and chemokines are produced by epidermal γδ T cells upon TCR signaling, we activated epidermal γδ T cells with anti-CD3 and evaluated cytokines and chemokines by Luminex assay. Epidermal γδ T cells produce a wide variety of cytokines and chemokines upon anti-CD3 stimulation (Fig. 3A). Cytokines and chemokines that exhibit over a 10-fold increase for all three cell lines tested include IL-2, IP-10 (CXCL10), IL-4, IL-13, IFN-γ, GM-CSF, RANTES (CCL5), MIP1a (CCL3), IL-17A, and MIP1β (CCL4). Many of these cytokines are associated with inflammation and cytokine storm (33,34). The identification of Th1, Th2, and Th17 cytokines and chemokines suggests that different functional subsets exist within epidermal γδ T cells. While direct TCR signals activate all epidermal T cell subsets, other stimuli such as TNF-α may stimulate particular functional subsets.

**Figure 3.**
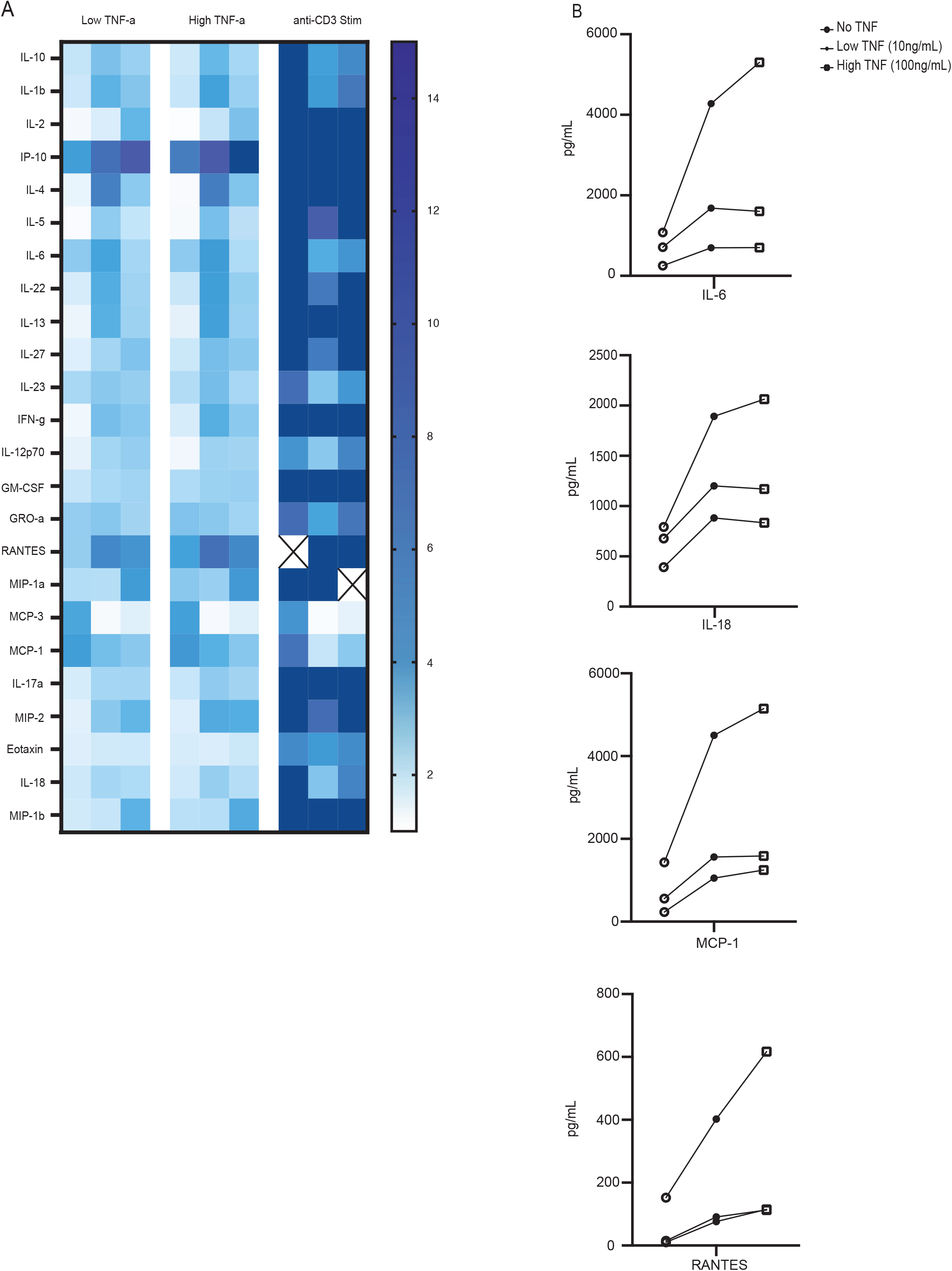
Epidermal γδ T cells produce IL-17 signature cytokines and chemokines in response to TNF-α treatment. (A) Heatmap showing relative fold change of inflammatory cytokine and chemokine production by epidermal γδ T cell lines stimulated with or with anti-CD3 and TNF-α. (B) Line graphs representing IL-6, IL-18, MCP-1, and RANTES concentrations with or without TNF-α addition. n = 3

Since a subset of epidermal γδ T cells expresses TNFR2, we evaluated whether low (10 ng/ml) or high (100 ng/ml) concentrations of TNF-α alone induces a distinct set of cytokines and chemokines. TNF-α induces epidermal γδ T cells to produce IL-17-associated cytokines and chemokines (above 2-fold change for all three cell lines): IL-1β, CXCL10, IL-6, IL-23, Gro-α (CXCL1), CCL5, CCL3, MCP1 (CCL2), and MIP2 (CXCL2). IL-1β and IL-6 are Th17 polarizing cytokines, IL-23 participates in IL-17A and F coproduction, CXCL1, CXCL2 and CCL2 are induced by IL-17. In addition, 2 out of 3 cell lines show a greater than 2-fold increase in IL-17A and IL22 (Fig. 3A). This shift from Th1, Th2, Th17 pleiotropic function in response to anti-CD3 to Th17-centric function with TNF-α stimulation suggests a role for TNF-α in inducing a specific subset of TNF-responsive epidermal γδ T cells.

### TNFR1 and TNFR2 define discrete subsets of epidermal γδ T cells with distinct functions

To identify the function of TNFR2-expressing epidermal γδ T cell subsets, we reanalyzed publicly available scRNAseq data from Liu et al, concentrating on TNFR1^+^ or TNFR2^+^ epidermal γδ T cell populations. The researchers treated C57BL/6J mice with or without 5% IMQ for a period of 7 days to induce psoriasis-like inflammation and isolated CD45^+^ skin cells for scRNA sequencing. We specifically analyzed epidermal γδ T cells using Loupe Browser and specific gene markers (*Cd3*^*+*^, *Tcrg-v5*^*+*^, *Fcer1g*^*+*^, *Cd4*^*−*^, *Cd8*^*−*^). Cells expressing *Tcrg-v4* and *Tcrg-v6* were also excluded. The result was 236 epidermal γδ T cells left for analysis (Fig. 4A). As we previously reported, epidermal γδ T cells displayed on a t-distributed stochastic neighbor embedding (t-SNE) plot define three subsets (clusters 1-3) with unique gene expression (Fig.4A)(35). The epidermal γδ T cells were further aggregated based on TNFR1 or 2 expression and treatment conditions (Fig. 4B). There are very few epidermal γδ T cells that express TNFR1 without TNFR2, defining a subset of TNFR1/2 double positive cells (Fig. 4B). Most TNFR1/2 double positive epidermal γδ T cells are found in cluster 1 before and after IMQ treatment suggesting they have functional homogeneity (Fig. 4B). There is also a population of epidermal

**Figure 4.**
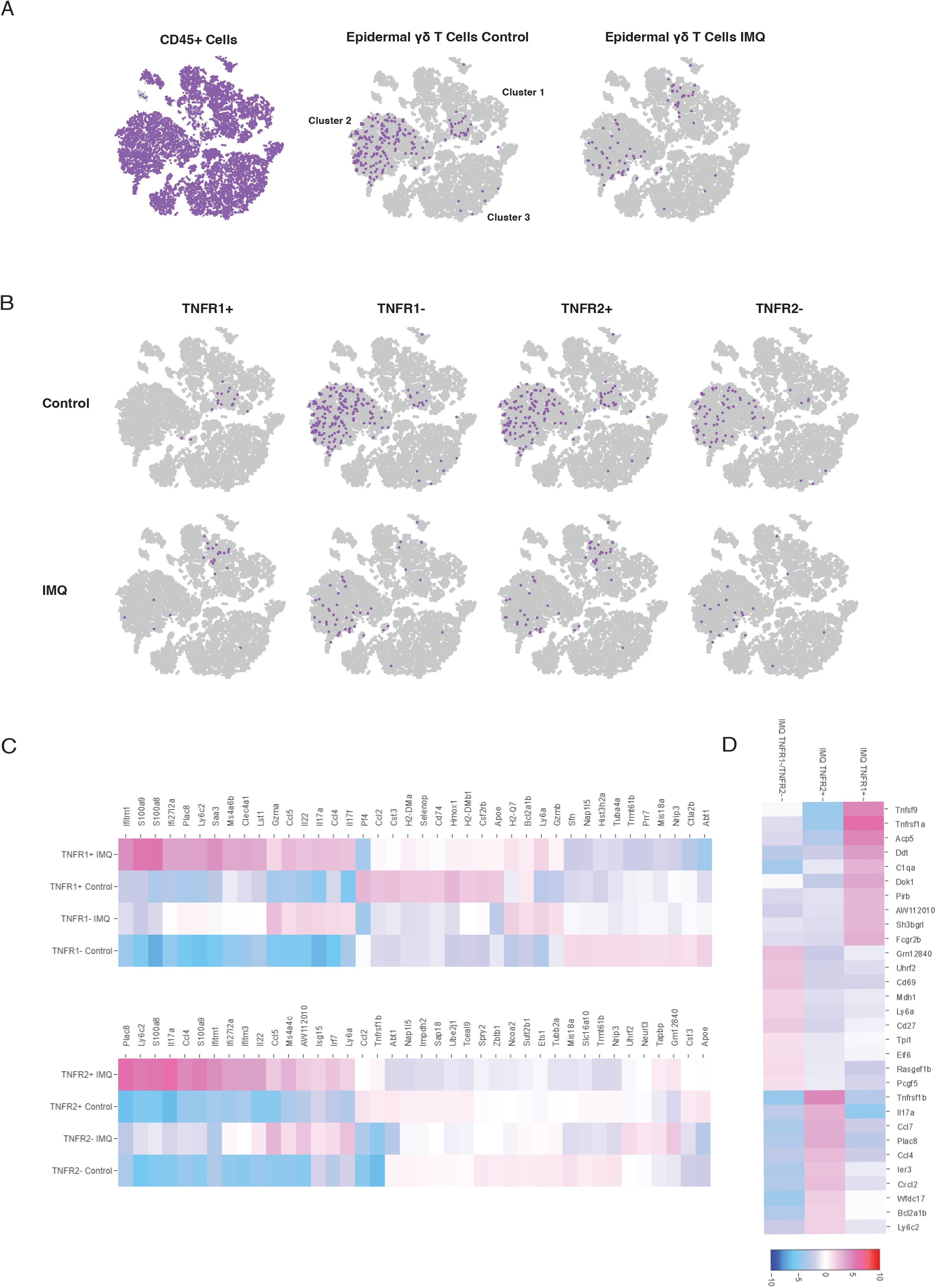
Epidermal γδ T cells expressing TNFR1 or TNFR2 present distinct gene expression profiles. (A) Publicly available scRNA-Seq data (Liu, Y. et al. 2020) total skin cell population (n=18,040) was filtered to only epidermal γδ T cells (n=236), then split between IMQ treatment and control groups. (B) UMAP plots showing filtered epidermal γδ T cells split into TNFR1+ and TNFR1-, TNFR2+ and TNFR2-subsets in IMQ treatment and control groups. (C) Heatmap renderings of differential gene expression between all epidermal γδ T cell treatments, as well as differential gene expression between control TNFR1+ and TNFR1-, TNFR2+ and TNFR2-, and IMQ treatment TNFR1+ vs. TNFR2+ vs. TNFR1-/ TNFR2-.

γδ T cells that only express TNFR2 and are found in all three clusters suggesting heterogeneity. Last, there is a population of epidermal γδ T cells that do not express TNFR1 or 2 that are also found in all three clusters suggesting heterogeneity within this population. Thus, epidermal γδ T cells in this dataset can be defined as TNFR negative, TNFR2 positive, or TNFR1/2 double positive.

A heat map displaying the top differentially expressed genes (DEGs) within TNFR1 or TNFR2-expressing γδ T cells was compared between untreated and IMQ treated mice (Fig. 4C). During IMQ treatment, TNFR1-expressing epidermal γδ T cells, which are mostly TNFR1/2 double positive, upregulate interferon-response genes (ifitm1, ifi2712a) and inflammatory response genes (s100a8, s100a9) as compared to TNFR1^+^ cells from untreated mice or TNFR1^-^ cells. TNFR1-expressing epidermal γδ T cells also exhibit a Th17 signature (*IL-22, CCL4, CCL5*, and *IL-17A)* although this is expressed at a lower level. TNFR2^+^ epidermal γδ T cells also exhibit a Th17 gene signature *IL-17A, IL-22, CCL4*, and *CCL5;* however, these are among the highest upregulated genes (Fig. 4C). This correlates with our findings by Luminex showing that epidermal γδ T cells exhibit a Th17 gene signature in response to TNF-α (Fig. 3). TNFR2-expressing epidermal γδ T cells also upregulate DEGs that include interferon-response genes (ifitm1, ifi2712a) and inflammatory responses (s100a8, s100a9) as compared to cells from untreated mice. This suggests that epidermal γδ T cells expressing both TNFR1 and 2 have both distinct and shared functions with TNFR2-expressing epidermal γδ T cells.

To further examine the distinct functions of epidermal γδ T cells from IMQ-treated mice, we compared the DEGs in TNFR1^+^ versus TNFR2^+^ versus TNFR negative-populations (Fig. 4D). TNFR1^+^ epidermal γδ T cells upregulate genes associated with negative regulation of T cell proliferation and activation (*Dok1, TNFsf9, Fcgr2b. Pirb)* when compared with cells that do not express TNFR1 or 2 or only express TNFR2. This suggests that the activated epidermal γδ T cells that express both TNFR1 and 2 upregulate receptors that control their proliferation and activation possibly in an effort to dampen inflammation. Once again, the TNFR2^+^ epidermal γδ T cells from IMQ mice express a Th17 inflammatory gene profile (*Il17a, Ccl7, Ccl4, Cxcl2*). In contrast, epidermal γδ T cells that lack TNFR1 and 2 express genes associated with activation (*Cd69* and *Ly6a*) and IFN-γ production (*CD27*). Thus, TNFR1 and/or 2 expression defines distinct functional epidermal γδ T cell populations during inflammatory skin disease.

### TNFR1 and TNFR2 epidermal γδ T cell subsets display an inflammatory gene profile in IMQ-treated mice

To gain unbiased insight into what distinguishes TNFR-expressing epidermal γδ T cell subsets during psoriasis, IPA core analysis was carried out to analyze biological and molecular pathways. The differentially expressed genes were clustered into canonical pathways by IPA Knowledge Base platform (Fig. 5). The comparison of TNFR1^+^ control to TNFR1^+^ IMQ populations identified reduced TCR Signaling pathways which correlate with TNFR1-expressing epidermal γδ T cells becoming overstimulated during skin inflammation and downregulating TCR signaling. The top pathway that is increased is Neutrophil Degranulation, which correlates with neutrophil attracting chemokines that are upregulated in the Luminex data (Fig. 3) suggesting a regulatory role for epidermal γδ T cells. The comparison of TNFR2^+^ control to TNFR2^+^ IMQ populations identifies Interferon alpha/beta signaling, Role of Hypercytokinemia/Hyperchemokemia in the Pathogenesis of Influenza, Differential Regulation of Cytokine Production in Macrophages and T Helper Cells by IL-17A and IL-17F, Differential Regulation of Cytokine Production in Intestinal Epithelial Cells by IL-17A and IL-17F, Neutrophil degranulation, Pathogen Induced Cytokine Storm Signaling Pathway, Interferon Signaling, and IL-23 Signaling Pathway to be upregulated in TNFR2+IMQ subsets in psoriasis. This further establishes TNFR2^+^ epidermal γδ T cells as IL-17 effectors during skin inflammation, while TNFR1^+^ epidermal γδ T cells mediate neutrophil infiltration and function.

**Figure 5.**
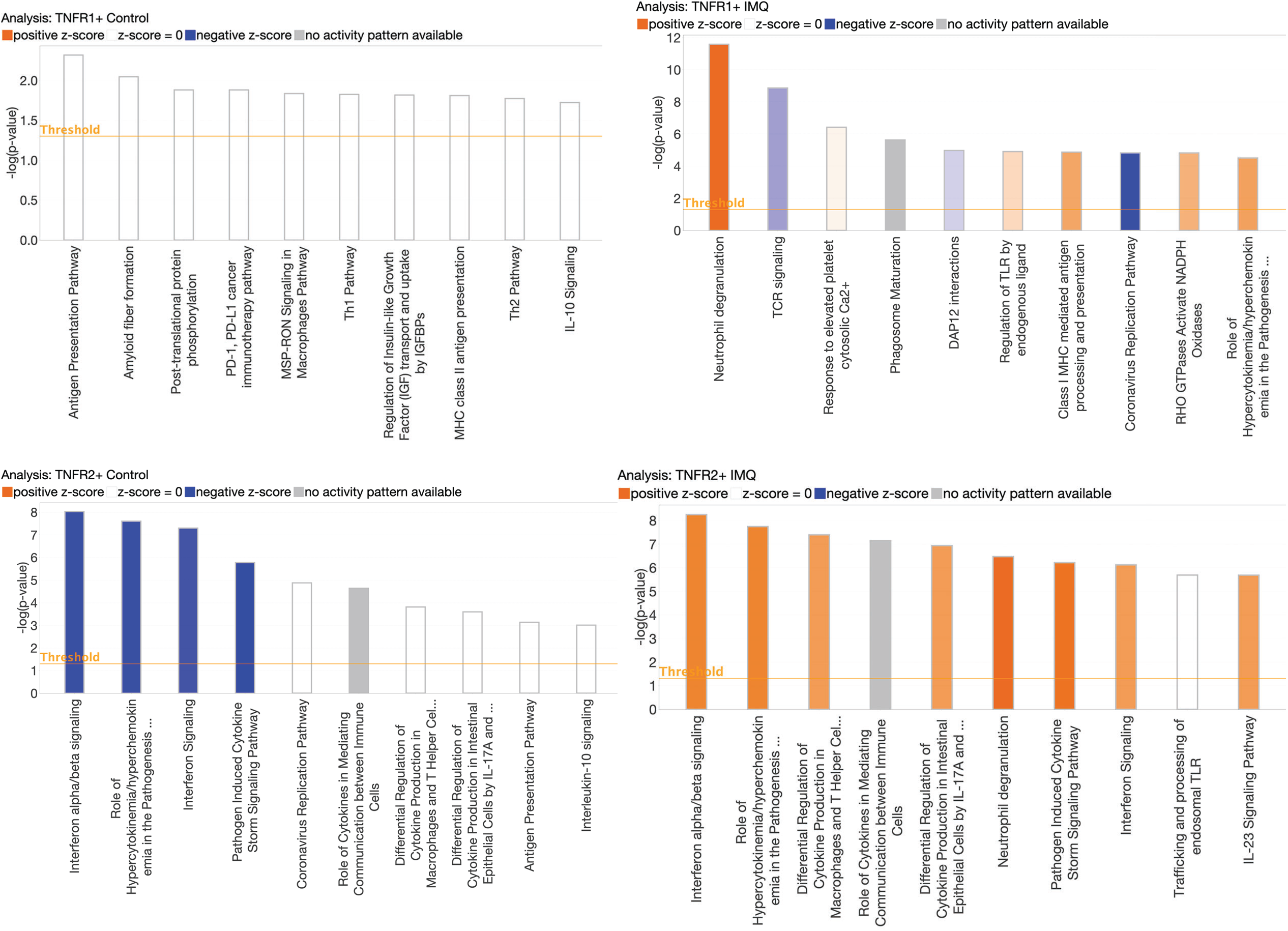
Canonical pathway analysis of differentially expressed genes from TNFR2^+^ epidermal γδ T cells during IMQ-induced psoriasis. The top 10 differentially expressed genes between TNFR2+IMQ and TNFR2+ control from publicly available scRNA sequencing were clustered in canonical pathways using the IPA knowledge Base platform. Publicly available scRNA sequencing data from Liu et al. A positive z score (orange) means the pathway is activated. A negative z-score means the pathway is inhibited. No z-score (white) means the pathway is not inhibited or activated.

## Discussion

Epidermal γδ T cells express a canonical Vγ5Vδ1 TCR that recognizes stressed, damaged, or transformed keratinocytes with the capacity to perform a range of functions in wound healing contact hypersensitivity, and cancer (18,25,36). Our findings suggest that there are distinct subsets among the population of epidermal γδ T cells that express TNFR2 and exhibit Tγδ17 functionality. This would imply a lack of uniformity within populations of epidermal γδ T cells and suggest that this heterogeneity is either due to different programming in the thymus or through differential environmental signals, like costimulation molecules or cytokines, within the skin (26,37). A subset of CCR6^+^ epidermal γδ T cells has recently been identified with Tγδ17 function (35). Here we demonstrate that this IL-17-producing epidermal γδ T cell subset is regulated by TNF-α through TNFR2. This provides further evidence that epidermal γδ T cells produce factors that exacerbate skin inflammation in diseases such as psoriasis.

TNF-α is a known regulator of T cell differentiation and IL-17 production (35,38). TNF-α induces the differentiation of CD4^+^ Th17 cells via TNFR2 signaling, resulting in heightened inflammation in EAE (38). In skin this contributes to inflammation by reinforcing the TNF-α/IL-17/IL-23 axis at multiple points due to the downstream effects in cellular targets such as keratinocytes and dendritic cells. (6,39) TNF-α can act with IL-17 and IL-22 to stimulate keratinocytes, which leads to the production of proinflammatory cytokines that contribute to the chronic activation of dendritic cells (6). Our findings reveal that TNFR2 is also expressed on a subset of epidermal γδ T cells in psoriasis with a Th17 gene profile and TNF-α induces Th17 signature cytokine and chemokine production. When considering the TNF/IL-23/IL-17 axis in psoriasis, our findings highlight a potential mechanism through which TNF-α amplifies inflammation via epidermal γδ T cells, contributing to the proinflammatory feedback loop in psoriasis (6).

During psoriasis we have found that epidermal γδ T cells are either TNFR1/2 negative, TNFR1/2 double positive or TNFR2 positive. While TNFR2-expressing cells express signature Tγδ17 genes, the additional upregulation of TNFR1 to a double positive cell dampens Tγδ17 gene expression and upregulates negative regulators of cell proliferation. This change in functionality can be indicative of a dynamic crosstalk between the two receptors. TNFR1 activation can influence TNFR2 signaling by regulating shared proteins, such as TRAF1 and TRAF2, thereby modifying TNFR2-controlled outcomes (40,41). We hypothesize that as inflammation persists, excess amounts of stimulation via TNF-α, may induce a subset of TNFR2^+^ epidermal γδ T cells to upregulate TNFR1 and dampen the response. This shift may serve as a mechanism to help combat chronic inflammation, especially within the context of psoriasis.

Treatment for psoriasis is partially dependent on the severity of the disease (42). In more severe cases, biologics that target the TNF-α/IL-23/IL-17 axis have enhanced efficacy during treatment (42). TNF-a inhibitors like etanercept work by lowering the level of cytokines produced by inflammatory dendritic cells. Skin resident γδ T cells are a likely target of the approved biologics for psoriasis due to their ability to produce IL-17 and contribute to the TNF-α/IL-23/IL-17 axis (43). Further investigation into how these resident cells respond to treatment could help to further refine and identify biological targets for future therapies.

## Acknowledgements

Thank you to the Jameson Lab for reviewing this manuscript.

## Abbreviations

IMQ: imiquimod
HFD: high fat diet
NCD: normal control diet
IP-10: Interferon Gamma-induced Protein 10
RANTES: Regulated on Activation, Normal T Expressed and Secreted
MIP1a: Macrophage Inflammatory Protein 1-alpha
MIP1b: Macrophage Inflammatory Protein 1 Beta
Gro-a: Growth-Related Protein-Alpha
MCP1: Monocyte Chemoattractant Protein-1
MIP2: Macrophage inflammatory protein 2

## References

1. Organisation mondiale de la santé, editor. Global report on psoriasis 2016. Geneva: World health organization; 2016.

2. Armstrong AW, Mehta MD, Schupp CW, Gondo GC, Bell SJ, Griffiths CEM. Psoriasis Prevalence in Adults in the United States. JAMA Dermatology. 2021 Aug 1;157(8):940–6.

3. Stuart PE, Nair RP, Ellinghaus E, Ding J, Tejasvi T, Gudjonsson JE, et al. Genome-wide association analysis identifies three psoriasis susceptibility loci. Nat Genet. 2010 Nov;42(11):1000–4.

4. Chandran V, Raychaudhuri SP. Geoepidemiology and environmental factors of psoriasis and psoriatic arthritis. Journal of Autoimmunity. 2010 May 1;34(3):J314–21.

5. Kim J, Moreno A, Krueger JG. The imbalance between Type 17 T-cells and regulatory immune cell subsets in psoriasis vulgaris. Front Immunol. 2022;13:1005115.

6. ten Bergen LL, Petrovic A, Krogh Aarebrot A, Appel S. The TNF/IL-23/IL-17 axis—Head-to-head trials comparing different biologics in psoriasis treatment. Scandinavian Journal of Immunology. 2020;92(4):e12946.

7. Nestle FO, Conrad C, Tun-Kyi A, Homey B, Gombert M, Boyman O, et al. Plasmacytoid predendritic cells initiate psoriasis through interferon-α production. Journal of Experimental Medicine. 2005 Jul 5;202(1):135–43.

8. Ganguly D, Chamilos G, Lande R, Gregorio J, Meller S, Facchinetti V, et al. Self-RNA– antimicrobial peptide complexes activate human dendritic cells through TLR7 and TLR8. J Exp Med. 2009 Aug 31;206(9):1983–94.

9. Res PCM, Piskin G, Boer OJ de, Loos CM van der, Teeling P, Bos JD, et al. Overrepresentation of IL-17A and IL-22 Producing CD8 T Cells in Lesional Skin Suggests Their Involvement in the Pathogenesis of Psoriasis. PLOS ONE. 2010 Nov 24;5(11):e14108.

10. Cai Y, Shen X, Ding C, Qi C, Li K, Li X, et al. Pivotal Role of Dermal IL-17-producing γδ T Cells in Skin Inflammation. Immunity. 2011 Oct 28;35(4):596–610.

11. Krueger JG, Brunner PM. Interleukin-17 alters the biology of many cell types involved in the genesis of psoriasis, systemic inflammation and associated comorbidities. Exp Dermatol. 2018 Feb;27(2):115–23.

12. Wang CQF, Akalu YT, Suarez-Farinas M, Gonzalez J, Mitsui H, Lowes MA, et al. IL-17 and TNF synergistically modulate cytokine expression while suppressing melanogenesis: potential relevance to psoriasis. J Invest Dermatol. 2013 Dec;133(12):2741–52.

13. Chiricozzi A, Guttman-Yassky E, Suárez-Fariñas M, Nograles KE, Tian S, Cardinale I, et al. Integrative Responses to IL-17 and TNF-α in Human Keratinocytes Account for Key Inflammatory Pathogenic Circuits in Psoriasis. Journal of Investigative Dermatology. 2011 Mar 1;131(3):677–87.

14. Zhou J, Zhang J, Tao L, Peng K, Zhang Q, Yan K, et al. Up-regulation of BTN3A1 on CD14+ cells promotes V?9Vd2 T cell activation in psoriasis. Proceedings of the National Academy of Sciences. 2022 Nov;119(44):e2117523119.

15. Boyman O, Hefti HP, Conrad C, Nickoloff BJ, Suter M, Nestle FO. Spontaneous Development of Psoriasis in a New Animal Model Shows an Essential Role for Resident T Cells and Tumor Necrosis Factor-α. Journal of Experimental Medicine. 2004 Feb 23;199(5):731–6.

16. Ruda RC, Kelly KA, Feldman SR. Real-world outcomes following switching from anti-TNF reference products to biosimilars for the treatment of psoriasis. Journal of Dermatological Treatment. 2023 Dec 31;34(1):2140569.

17. Lawler W, Castellanos T, Engel E, Alvizo CR, Kasler A, Bshara-Corson S, et al. Impact of obesity on the CCR6-CCL20 axis in epidermal γδ T cells and IL-17A production in murine wound healing and psoriasis. J Immunol. 2025 Jan 1;214(1):153–66.

18. Jameson J, Ugarte K, Chen N, Yachi P, Fuchs E, Boismenu R, et al. A role for skin gammadelta T cells in wound repair. Science. 2002 Apr 26;296(5568):747–9.

19. Sharp LL, Jameson JM, Cauvi G, Havran WL. Dendritic epidermal T cells regulate skin homeostasis through local production of insulin-like growth factor 1. Nat Immunol. 2005 Jan;6(1):73–9.

20. Toulon A, Breton L, Taylor KR, Tenenhaus M, Bhavsar D, Lanigan C, et al. A role for human skin–resident T cells in wound healing. Journal of Experimental Medicine. 2009 Mar 23;206(4):743–50.

21. Heilig JS, Tonegawa S. Diversity of murine gamma genes and expression in fetal and adult T lymphocytes. Nature. 1986 Sep 28;322(6082):836–40.

22. Havran WL, Allison JP. Origin of Thy-1+ dendritic epidermal cells of adult mice from fetal thymic precursors. Nature. 1990 Mar;344(6261):68–70.

23. Havran WL, Chien YH, Allison JP. Recognition of Self Antigens by Skin-Derived T Cells with Invariant γδ Antigen Receptors. Science. 1991 Jun 7;252(5011):1430–2.

24. Jameson JM, Cauvi G, Sharp LL, Witherden DA, Havran WL. γδ T cell–induced hyaluronan production by epithelial cells regulates inflammation. J Exp Med. 2005 Apr 18;201(8):1269–79.

25. Girardi M, Glusac E, Filler RB, Roberts SJ, Propperova I, Lewis J, et al. The distinct contributions of murine T cell receptor (TCR)gammadelta+ and TCRalphabeta+ T cells to different stages of chemically induced skin cancer. J Exp Med. 2003 Sep 1;198(5):747–55.

26. Witherden DA, Verdino P, Rieder SE, Garijo O, Mills RE, Teyton L, et al. The junctional adhesion molecule JAML is a costimulatory receptor for epithelial gammadelta T cell activation. Science. 2010 Sep 3;329(5996):1205–10.

27. Whang MI, Guerra N, Raulet DH. Costimulation of Dendritic Epidermal γδ T Cells by a New NKG2D Ligand Expressed Specifically in the Skin 1. The Journal of Immunology. 2009 Apr 15;182(8):4557–64.

28. Hsu H, Xiong J, Goeddel DV. The TNF receptor 1-associated protein TRADD signals cell death and NF-kappa B activation. Cell. 1995 May 19;81(4):495–504.

29. Chen S, Lin Z, Xi L, Zheng Y, Zhou Q, Chen X. Differential role of TNFR1 and TNFR2 in the development of imiquimod-induced mouse psoriasis. J Leukoc Biol. 2021 Dec;110(6):1047–55.

30. Chandrasekharan UM, Kaur R, Harvey JE, Braley C, Rai V, Lee M, et al. TNFR2 Depletion Reduces Psoriatic Inflammation in Mice by Downregulating Specific Dendritic Cell Populations in Lymph Nodes and Inhibiting IL-23/IL-17 Pathways. Journal of Investigative Dermatology. 2022 Aug 1;142(8):2159-2172.e9.

31. Jameson JM, Cauvi G, Witherden DA, Havran WL. A Keratinocyte-Responsive γδ TCR Is Necessary for Dendritic Epidermal T Cell Activation by Damaged Keratinocytes and Maintenance in the Epidermis1. The Journal of Immunology. 2004 Mar 15;172(6):3573–9.

32. Liu Y, Cook C, Sedgewick AJ, Zhang S, Fassett MS, Ricardo-Gonzalez RR, et al. Single-Cell Profiling Reveals Divergent, Globally Patterned Immune Responses in Murine Skin Inflammation. iScience. 2020 Sep 19;23(10):101582.

33. Karki R, Kanneganti TD. The ‘cytokine storm’: molecular mechanisms and therapeutic prospects. Trends in Immunology. 2021 Aug 1;42(8):681–705.

34. Coperchini F, Chiovato L, Croce L, Magri F, Rotondi M. The cytokine storm in COVID-19: An overview of the involvement of the chemokine/chemokine-receptor system. Cytokine & Growth Factor Reviews. 2020 Jun 1;53:25–32.

35. Lawler W, Castellanos T, Engel E, Alvizo CR, Kasler A, Bshara-Corson S, et al. Impact of obesity on the CCR6-CCL20 axis in epidermal γδ T cells and IL-17A production in murine wound healing and psoriasis. The Journal of Immunology. 2025 Jan 1;214(1):153–66.

36. Nielsen MM, Lovato P, MacLeod AS, Witherden DA, Skov L, Dyring-Andersen B, et al. IL-1ß–Dependent Activation of Dendritic Epidermal T Cells in Contact Hypersensitivity. The Journal of Immunology. 2014 Apr 1;192(7):2975–83.

37. Ribot JC, deBarros A, Pang DJ, Neves JF, Peperzak V, Roberts SJ, et al. CD27 is a thymic determinant of the balance between interferon-gamma- and interleukin 17-producing gammadelta T cell subsets. Nat Immunol. 2009 Apr;10(4):427–36.

38. Alam MS, Otsuka S, Wong N, Abbasi A, Gaida MM, Fan Y, et al. TNF plays a crucial role in inflammation by signaling via T cell TNFR2. Proceedings of the National Academy of Sciences. 2021 Dec 14;118(50):e2109972118.

39. Furue K, Ito T, Furue M. Differential efficacy of biologic treatments targeting the TNF-a/IL-23/IL-17 axis in psoriasis and psoriatic arthritis. Cytokine. 2018 Nov 1;111:182–8.

40. Naudé PJW, den Boer JA, Luiten PGM, Eisel ULM. Tumor necrosis factor receptor cross-talk. The FEBS Journal. 2011;278(6):888–98.

41. Wicovsky A, Henkler F, Salzmann S, Scheurich P, Kneitz C, Wajant H. Tumor necrosis factor receptor-associated factor-1 enhances proinflammatory TNF receptor-2 signaling and modifies TNFR1–TNFR2 cooperation. Oncogene. 2009 Apr;28(15):1769–81.

42. Lee HJ, Kim M. Challenges and Future Trends in the Treatment of Psoriasis. Int J Mol Sci. 2023 Aug 28;24(17):13313.

43. Qi C, Wang Y, Li P, Zhao J. Gamma Delta T Cells and Their Pathogenic Role in Psoriasis. Front Immunol. 2021 Feb 25;12:627139.

